# Chromosome-level genome assembly of the Northeast China Brown Frog (*Rana dybowskii*)

**DOI:** 10.64898/2026.06.11.731602

**Authors:** Yuanyuan Zhang, Di Wang, Ran Zhao, Shaowu Li, Xianhu Zheng, Guo Hu

## Abstract

*Rana dybowskii* is distributed across Northeast Asian and represents a valuable medical resource. A high-quality assembly of the genome has not yet been reproted. This species has 2n=24 chromosomes, but a huge genome size that estimated at 3.5 ~4.6 Gb in the previous studies. The relatively large chromosome size, exceeding hundreds of megabases, may result in difficulties of obtaining a complete chromosome level genome. Here, we constructed a chromosome-level genome assembly of *R. dybowskii* by integrating PacBio HiFi long-read sequencing for de novo assembly and CiFi (3C coupled with HiFi sequencing) for scaffolding. The final assembly consists of 12 chromosomes with a total of 3.95 Gb and a scaffold N50 length of 455 Mb. BUSCO assessment using the tetrapoda_odb12 database identified 94.2% complete and 0.5% fragmented orthologs, suggesting a high level of completeness of the assembly. Genomic annotation revealed that repetitive sequences comprise over 53% of the assembly, with retroelements and DNA transposons accounting for 22% and 25%, respectively. A total of 43,999 protein-coding genes were predicted with the assistance of RNA-seq reads from four tissues (muscle, eye, testis and skin). This high-quality chromosome-level reference genome provides a valuable genomic resource for advancing genetic studies of the species.

## Background & Summary

*Rana dybowskii* belongs to the phylum Chordata, subphylum Vertebrata, class Amphibia, order Anura, and family *Ranidae*. It is predominantly distributed in Northeast China^1^, including Heilongjiang, Jilin, Liaoning, and eastern Inner Mongolia, and also can be habituated in Russia, North Korea, Mongolia and Japan. It has a long history of medicinal use, particularly the dried oviducts of females, which are widely applied in traditional Chinese medicine. Its medicinal value was first documented for medicine value since the Ming Dynasty as “Xue Ha (Snow frog)”in the *Compendium of Materia Medica (Bencao Gangmu)*. By the mid-to-late Qing Dynasty, under the influence of cultural integration between the Manchu and Han populations, it became widely known as “Hashima” in northern China and the dried oviduct of female forest frogs from Northeast China was referred as “Hamayou” (Ranae Oviductus) and officially documented included in the Chinese Pharmacopoeia. The 2025 edition specifies *Rana dybowskii Günther* as the authentic source species^2^. Several researches of Oviductus Ranae reported it could have medicinally valuable components that exhibits diverse biological activities, including antioxidant, anti-osteoporotic, antitumor, immunomodulatory, and hypolipidemic effects^3–5^.

In Northeast China, *R. dybowskii* plays a vital role in forest-wetland ecosystems and has been historically cultivated as an economically amphibian. Currently, the farming industry supports tens of thousands of households and practitioners, generating an annual output value of approximately 20 billion Chinese yuan, which underscoring its considerable economic and medicinal importance. Consequently, genetic breeding and biological analysis of *R. dybowskii* are quite emergency for guiding the efficient breeding and biological investigations of *R. dybowskii* are urgently required to guide efficient feeding and breeding programs and promote its biological and medical applications. However, genomic resources and functional gene information for this species remain limited. A previous study assembled the genome of a male individual using next-generation sequencing short reads, resulting in 270,785 contigs and 498 scaffolds, this assembled genome size was approximately 3.58 Gb, with a total of 40,913 genes predicted^6^. Although this assembly provided valuable genomic sequences and gene information, its continuity and accuracy were insufficient for further genomic and molecular mechanism analysis.

In the present study, we generated a high-quality, highly contiguous chromosome-scale genome of *R. dybowskii* using PacBio HiFi reads to generate high-quality contigs, and scaffolding using long reads of chromatin interaction produced with PacBio HiFi sequencing (CiFi). The final assembly spanned approximately 3.95 Gb, comprising 12 nuclear chromosomes and one mitochondrial sequence. The largest chromosome assembled to be over 604 Mb and the scaffold N50 reached 455 Mb. BUSCO assessment analysis based on the tetrapoda_odb12 dataset recovered nearly 95% complete orthologs, which represents a substantial improvement in genome continuity and completeness.

## Methods

### Sample collection

Adult individuals of *R. dybowskii* used in the present study was collected on December, 2025 from Daxilin Forest Farm, YiChun City, Heilongjiang Province, China (47.6361° N, 129.1334° E). Animals were humanely euthanized after being anesthetized by placing a cotton ball soaked with 99% ethanol into the mouth. Under anesthesia, the spinal cord was destroyed using scissors. All the tissue samples were immediately flash-frozen in liquid nitrogen and then stored at −80°C for the further using. A male individual (Figure 1a) was selected for genome assembly, genomic DNA was extracted from liver and muscle tissues and used for whole-genome resequencing, PacBio HiFi sequencing, and CiFi library construction. All the animal experiments were approved by the Institutional Review Board (IRB) of the Heilongjiang River Fisheries Research Institute for Laboratory Animal Welfare.

**Figure 1.**
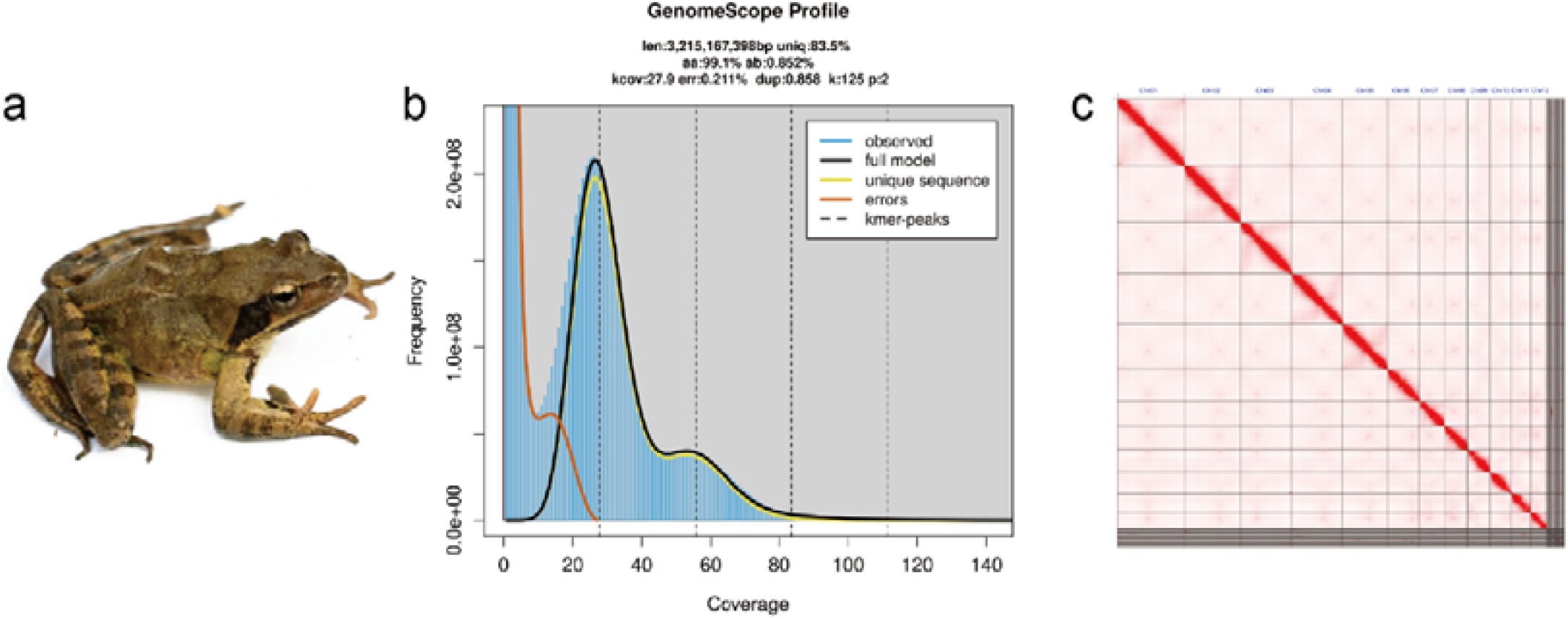
Genome survey and chromosomal scaffolding by CiFi reads. (a) A photograph of a male *R. dybowskii*, provided by Guo Hu. (b) Genome size estimated by GenomeScope using 125-mers calculated by KMC. (c) Chromosomal-level scaffolding of the *R. dybowskii* assembly based on alignments and interactions of the CiFi reads.

### Genomic and transcriptomic sequencing

DNA and RNA samples were constructed for sequencing libraries according to the manufacturer’s protocols for the further short reads sequencing on Illumina NovaSeq600 platform, the pacbio HiFi sequencing on Pacbio Revivo platform. The chromatin interaction sequencing with PacBio HiFi sequencing (CiFi) was prepared following a modified version of the protocol described by McGinty *et al*. ^7^ Briefly, fresh tissues were cross-linked with 1% formaldehyde, quenched with glycine, and lysed to release intact nuclei. Chromatin was digested using the restriction enzyme *DpnII*, followed by proximity ligation to generate long-range chromatin interaction products. Cross-links were subsequently reversed by proteinase K digestion, and long DNA was purified using phenol–chloroform extraction and ethanol precipitation. Size selection was performed using diluted AMPure PB beads prior to amplification. The purified 3C libraries were then amplified and converted into SMRTbell templates using the PacBio Ampli-Fi protocol. Final libraries were sequenced on the PacBio Revio platform to generate HiFi reads. Detailed sequenced data of all the samples summarized in table 1 and supplementary table 1–3.

**Table 1.** Summary of sequencing data in the present study.

| Library type | sample id | Clean data (Gb) | Read length (bp) | Coverage (×) |
| --- | --- | --- | --- | --- |
| WGS | R1 | 292.12 | 150 | 73.95 |
| WGS | R2 | 54.91 | 150 | 13.89 |
| RNA | R1E | 6.66 | 150 | - |
| RNA | R1G | 10.54 | 150 | - |
| RNA | R1M | 7.12 | 150 | - |
| RNA | R1S | 7.9 | 150 | - |
| HiFi | R1 | 359.76 | 18,704 (N50) | 91.08 |
| CiFi | R1 | 80.07 | 5,680 (N50) | - |

### Genome assessment and assembly

Single-molecule real-time circular consensus sequencing (CCS) library preparation was conducted following the manufacture’s protocol. A total of 238 Gb of HiFi reads generated by the PacBio Revio platform, corresponding to about 91 × genome coverage based on the estimated genome size. *k-mer* frequency was performed using KMC ^8^ (version 3.2.4) with a k-mer size of 125. Then GenomeScope2^9^ was subsequently applied to assess the genome size, heterozygosity and repetitive sequences content (Figure 1b). The error rate of HiFi reads is about 0.21 % at k=125, the long and accurate reads allow an accurate estimate of the genome size when using a large k-mer size, the genome estimated to be 3.22 Gb, these high-quality reads provided a solid foundation for accurate assembly and phasing of the diploid genome (Fig. 1b).

The primary genome assembly was produced by hifiasm ^10^ (version 0.25.0-r726) using HiFi reads longer than 1kb, yielding 1,063 contigs with a contig N50 of 46.17 Mb and a total assembly size of approximately 4.06 Gb (Table 2). Despite the high continuity achieved at the contig level, chromosome-scale assembly of *Rana dybowskii* remains challenging due to its ultra-long chromosomes exceeding several hundred megabases.

**Table 2.** Statistics of assembled genome.

|  |  |
| --- | --- |
| Total Length (bp) | 3,951,792,017 |
| Count of contigs | 1,063 |
| Contig N50 (bp) | 46,165,982 |
| Contig N90 (bp) | 3,594,215 |
| Count of scaffolds | 84 |
| Contig N50 (bp) | 455,097,160 |
| Contig N90 (bp) | 191,900,750 |
| Max chromosome length (bp) | 604,322,790 |
| Chromosome number | 12 |
| Chromosome length (bp) | 3,796,122,376 |
| Mitochondria length (bp) | 17,541 |
| Anchored rate (%) | 96.06 |
| Count of transcripts | 43,999 |
| BUSCO evaluation (tetrapoda_odb12) | C:94.2% [S:90.2%, D:4.0%], F:0.5%, M:5.3%, n:5623 |

To obtain ultra-long and highly continuous chromosome-scale scaffolds spanning hundreds of metabases employed CiFi reads for chromosome-scale scaffolding. CiFi technology integrates the advantages of long-range chromatin interaction information and high base-level accuracy derived from 3C-based proximity ligation, and has demonstrated excellent performance in large-scale genome scaffolding ^11^. Briefly, the CiFi reads were firstly processed with the Pore-C Nextflow pipeline ^12^ with minor modifications in the alignment step, in which minimap2 used for PacBio HiFi reads mapping using the “-x map-hifi” preset. CiFi segments within each HiFi read were then converted into a mock Illumina-like paired-end alignments bam file, representing all possible intramolecular chromatin interactions. PairTools^13^ (version 1.1.3) was applied to generate contact pairs, which were subsequently used by YAHS^14^ (version 1.2.2) for chromosome-scale scaffolding. This CiFi assistant scaffolding strategy enabled the successful reconstruction of ultra-long chromosomes, yielded twelve chromosomes with a highly contiguous assembly of 4.05 Gb (Figure 1c). Notably, the largest scaffold is over 600 Mb and all the pseudochromosomes were longer than 150 Mb, reflecting the exceptional continuity achieved for this large and complex amphibian genome. Purge_dups ^15^ (version 1.2.6) was used to remove duplicated haplotigs and heterozygous overlaps in the assembly. TGS-gapcloser^16^ (version 1.2.1) was used to close assembly gaps with parameters –min_match 1000–min_nread 3” using long HiFi reads that longer than 10 kb. Then the assembly was further polished using NextPolish2 ^17^ (version 0.2.2) with the whole genomic resequencing Illumina PE reads and PacBio HiFi reads larger than 1 kb to correct SNV and Indel errors avoiding overcorrections in regions rich of complex repeat elements. The final assembly spanned approximately 3.95 Gb, comprising 12 chromosome-scale pseudomolecules and 72 unplaced sequences, with the largest pseudochromosome reaching 604.32 Mb and an overall scaffold N50 of 455.01 Mb, and all the pseudochromosome larger than 150 Mb (Figure 2, Table 2 and 3).

**Table 3.** Chromosome lengths and features.

| Chromosome | Length (bp) | GC content (%) | Genes |
| --- | --- | --- | --- |
| Chr01 | 604,322,790 | 43.18 | 4,930 |
| Chr02 | 509,327,144 | 43.29 | 4,731 |
| Chr03 | 464,717,231 | 43.48 | 4,747 |
| Chr04 | 455,097,160 | 43.51 | 4,234 |
| Chr05 | 402,975,683 | 43.44 | 3,686 |
| Chr06 | 298,159,558 | 44.4 | 3,793 |
| Chr07 | 221,860,801 | 43.92 | 2,594 |
| Chr08 | 213,162,874 | 44.15 | 2,463 |
| Chr09 | 202,537,032 | 43.99 | 2,685 |
| Chr10 | 191,900,750 | 44.56 | 3,141 |
| Chr11 | 169,829,751 | 44.05 | 2,106 |
| Chr12 | 153,435,866 | 45.14 | 2,272 |
| MT | 23,402 | 39.3 | 37 |

**Figure 2.**
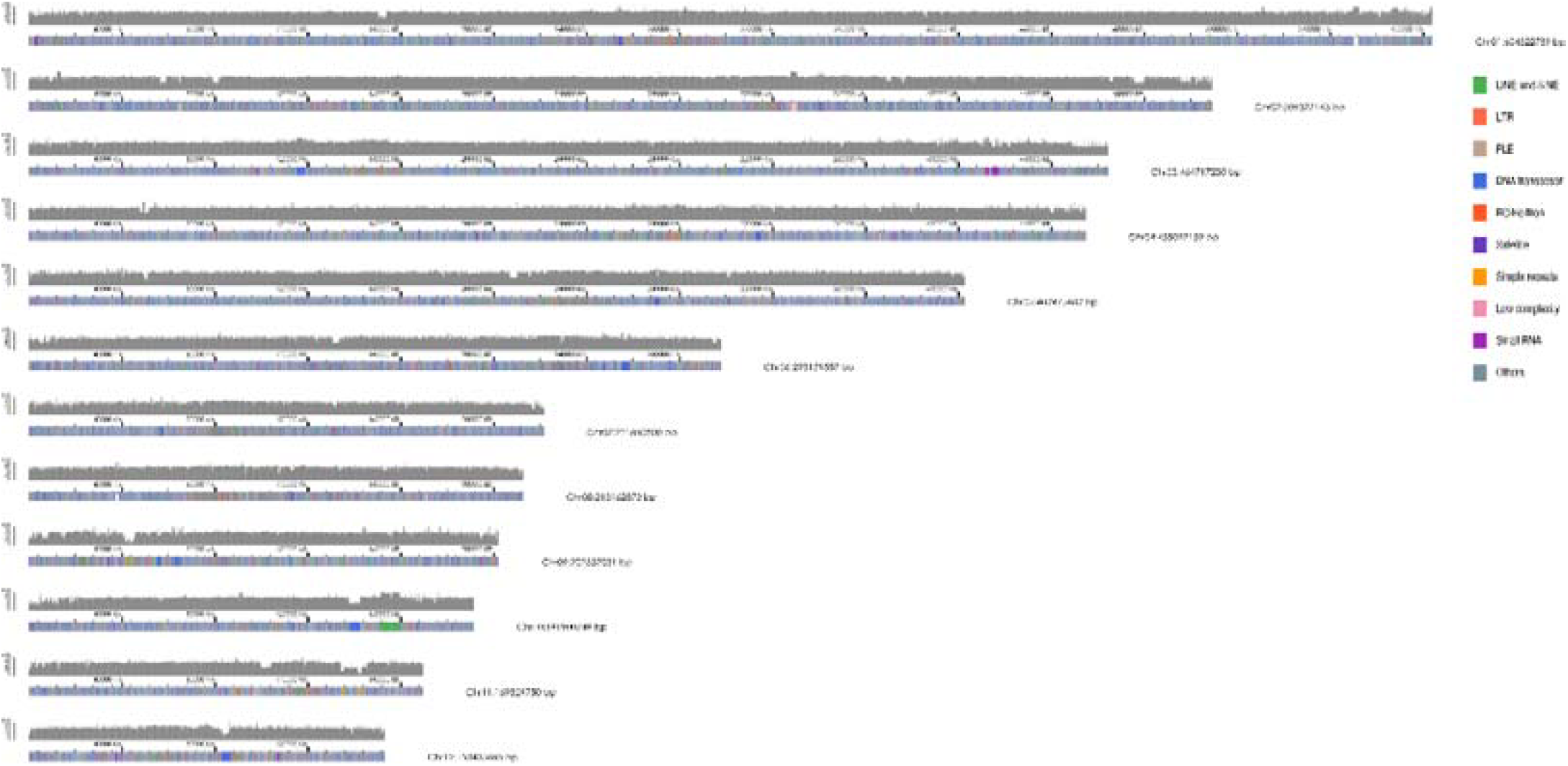
The assembled twelve chromosomes and resequencing coverage of a *R. dybowskii* male individual R1. The final genome assembly of R1 comprises twelve chromosomes. Whole-genome resequencing data (~80× coverage) were aligned to the reference genome, the calculated coverage depth within each 100kb window is displayed as grey line. Repeat elements, shown in different colors, are distributed across the entire genome.

### Mitochondrial sequence detection

A previous version of the mitochondrion (MT) sequence of *R. dybowskii* has been deposited in NCBI genebank with accession no. NC_023528.1, with a total length of 18,864 bp ^18^. During the assembly process, a contig containing more than 4 copies of the full length of the MT was identified using blastn^19^. Based on sequence alignment and boundary confirmation, a single complete mitochondrial genome of *R. dybowskii* was subsequently extracted from this contig using MitoHiFi ^20^ (version 3.2.1), yielding a mitochondrial sequence of 21,198 bp, including 37 genes (22 t-RNAs, 2 rRNAs and 13 protein-coding genes).

### Telomere and TE detection

Telomere repeats (AACCCT) were detected at one end of the pseudochromosome 1,3,11 and 12, as well as at one end of eight other unplaced scaffolds. This founding suggests the major telomere repeats is “AACCT”. However, these repeats were detected by HiFi sequencing but not successfully assembled into chromosomal ends, likely due to the challenges associated with assembling extremely long chromosomal regions. In the future study, ultra-long sequencing data, such as nanopore reads, could be involved and help improve telomere-to-telomere continuity in the next version of assembly. Nevertheless, the present assembly provides a high-quality reference for the genetic and biological studies of *R. dybrowskii*.

Repetitive elements in the final assembly were annotated using Repeatmodeler ^21^ and RepeatMasker ^22^ (version 4.09 with RepBase 20181026 and FamDB), and also searching using EDTA ^23^ pipeline in order to scan for more unclassified elements. RepeatMasker identified over 74% of the genome as repetitive elements by using “-uncurated” parameter, while the EDTA annotated 51.48% Class I and ClassII repetitive element. All the outputs were combined and finally identified over 53.85% of the genome sequences are repetitive sequences (Table 4), and visualized on the chromosomes (Figure 2).

**Table 4.** Summary of repetitive elements in the assembly.

| Class/Subclass | Number of elements | Total length (bp) | % of genome |
| --- | --- | --- | --- |
| **Class I (Retrotransposons)** | **2,111,071** | **873,830,307** | **22.11** |
| **LINE** | **865,780** | **408,828,736** | **10.35** |
| CR1 | 192,864 | 72,697,291 | 1.84 |
| L1 | 376,318 | 210,000,023 | 5.31 |
| L2 | 232,973 | 104,326,159 | 2.64 |
| R1 | 362 | 271,099 | 0.01 |
| R2 | 12,258 | 4,630,949 | 0.12 |
| RTE | 5,006 | 3,384,510 | 0.09 |
| Rex | 554 | 153,603 | <0.01 |
| unknown | 45,445 | 13,365,102 | 0.34 |
| **LTR** | **1,110,448** | **410,345,642** | **10.38** |
| Bel_Pao | 18,618 | 7,040,620 | 0.18 |
| Copia | 5,338 | 2,090,920 | 0.05 |
| Gypsy | 112,315 | 65,521,603 | 1.66 |
| Retrovirus | 3,416 | 892,408 | 0.02 |
| unknown | 970,761 | 334,800,091 | 8.47 |
| **SINE** | **54,622** | **21,090,717** | **0.53** |
| **nonLTR (Penelope)** | **80,221** | **33,565,212** | **0.85** |
| **Class II (DNA Transposons)** | **3,782,683** | **1,025,037,412** | **25.94** |
| **TIR** | **2,786,699** | **772,578,496** | **19.55** |
| CACTA | 477,454 | 116,727,514 | 2.95 |
| Mutator | 251,390 | 65,219,650 | 1.65 |
| PIF_Harbinger | 685,220 | 202,668,030 | 5.13 |
| Tc1_Mariner | 24,526 | 6,222,649 | 0.16 |
| hAT | 1,348,109 | 381,740,653 | 9.66 |
| **nonTIR (Helitron)** | **995,984** | **252,458,916** | **6.39** |
| **Unclassified** | **449,771** | **152,345,244** | **3.86** |
| **Small RNA** | **47,240** | **8166295** | **0.21** |
| **Satellites** | **69,288** | **24930867** | **0.63** |
| **Simple repeats** | **612,922** | **37065946** | **0.94** |
| **Low complexity** | **78,957** | **6409168** | **0.16** |
| **Total** | **7,151,932** | **2,127,785,239** | **53.85** |

### Gene prediction and annotation

The RNA-seq data from four tissues (muscle, eye, liver and skin) were align against with the assembled genome using STAR ^24^ (version 2.7.10b) to obtain the cDNA match evidence, which then be used in gene prediction model using BRAKER3 ^25^. De novo transcripts assembly constructed using Trinity ^26^ (version 2.15.2) were aligned to the genome and processed using PASA^27^ (version 2.5.3) to refine gene structures. Then the pre-assembled unigenes, and the predicted genes from BRAKER3 were integrated using the EVidenceModeler (EVM)^28^ software(version 2.1.0) to combine predicted genes and transcript alignments into weighted consensus gene models and combined these exonic evidences into a single automated gene structure annotation system. Finally, 43,999 genes were obtained in the assembled genome (table 3). Functional annotation of these genes were performed by eggnog-mapper^29,30^ (version 2.1.12) diamond tool with eggNOG 5.0 database13.

### Genome synteny analysis

Synteny analysis was performed between our assembly of *R. dybowskii* with the *R. temporaria* (Genome assembly accession no. GCA_905171725.1). Homologous proteins were identified using blastp (-evlaue 1e-6) and syntenic dotplot was generated using WGDI software^31^.

The self-synteny analysis of *R. dybowskii* (Figure 3a) revealed overall collinearity across the twelve chromosomes, while secondary alignments were observed on the end of Chr03 and Chr10, which may reflect the presence of homologous or repetitive regions within the genome. Similar secondary alignments were also detected in the corresponding orthologous regions in the inter-species comparison between *R. dybowskii* and *R. temporaria* (Figure 3b). Chr06 of *R. dybowskii* was consistently syntenic with two chromosomes (Chr11 and Chr13) in *R. temporaria*, suggesting a chromosomal fusion event in the *R. dybowskii* lineage (Figure 3b).

**Figure 3.**
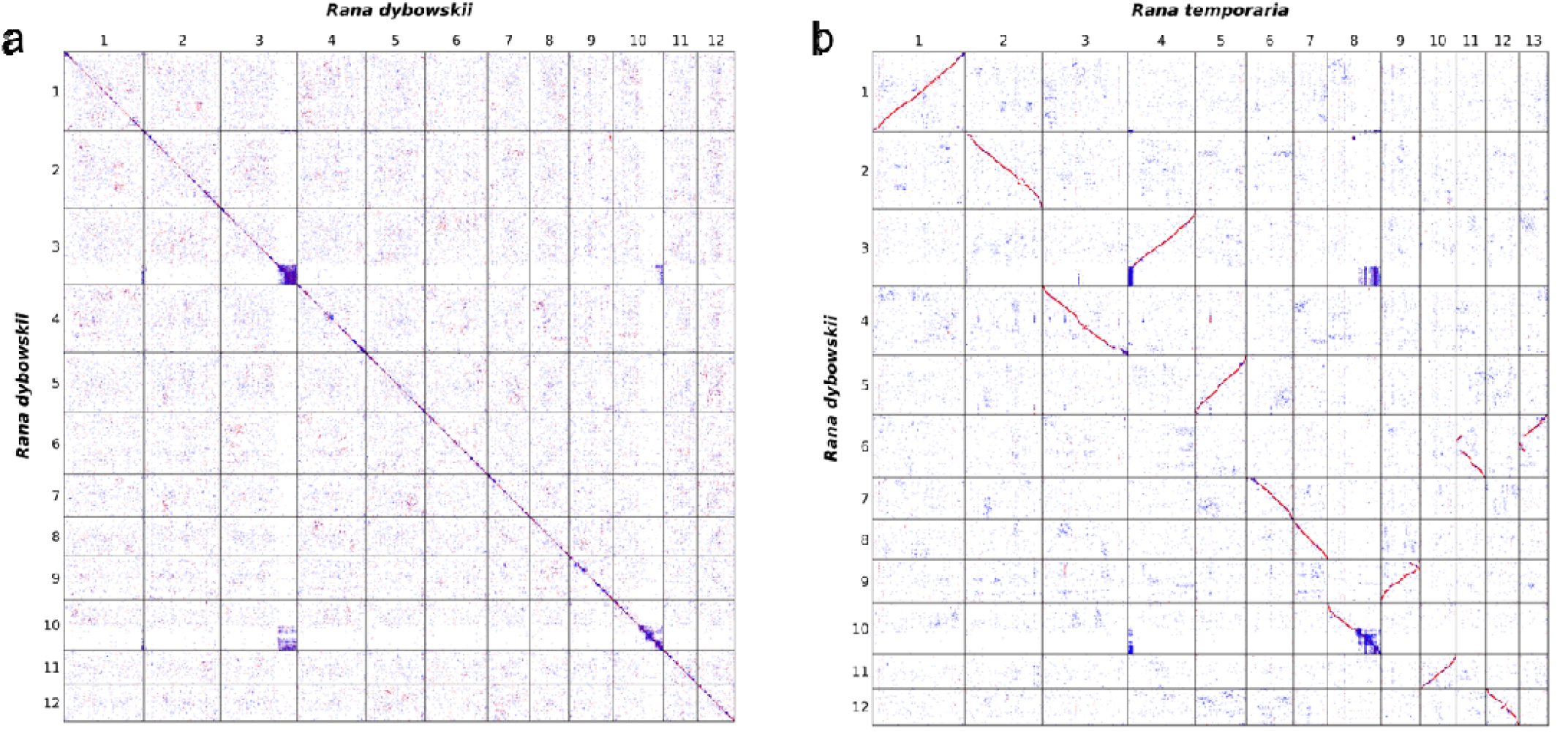
Syntenic dot plot analysis of *R. dybowskii* and *R. temporaria* genomes. (a) Self-alignment dot plot of the *R. dybowskii* genome, revealing intra-genomic synteny and clear repeat region on the end of chr03 and Chr10. (b) Cross-species syntenic dot plot between *R. dybowskii* and *R. temporaria*. The haploid chromosome number of *R. dybowskii* is 12, which is one less than that of *R. temporaria* (n = 13). Syntenic analysis suggests that Chr06 in *R. dybowskii* may have resulted from a fusion of two ancestral chromosomes corresponding to the two chromosomes (Chr11 and Chr13) in *R. temporaria*.

## Data Records

The sequenced reads reported in the present study have been deposited in China National Genomics Data Center (https://www.cncb.ac.cn/), and can be accessed through the project ID PRJCA051591. The genome assembly and annotation data has been deposited in the public database GWH under a preliminary accession no. <u>Batch0092431</u> and will be made publicly available upon publication of the manuscript.

## Technical Validation

The quality and concentration of DNA and RNA using 0.8 % agarose gel electrophoresis and Qubit 3.0 fluorometer (Life Technologies, Carlsbad, CA, USA). Nucleic acid samples with intact high-molecular-weight fragments and passed the quality control were used in the downstream library’s construction and sequencing.

No contaminants were found in the final assembled genome using the NCBI FCS tool suite^32^ (version 0.5.5). The quality control of genome The BUSCO^33,34^ (version 6.0.0) was used to assess the completeness and accuracy of the assembled genome using tetrapoda_odb12 database (n = 5623). For the BUSCO analysis, 94.2% of genes were completely recalled, with 90.2 % of them are single copies, and 4.0% and 0.5%of them originated from duplication and fragment events, respectively. Additionally, the NGS of the same individual used for assembly aligned against the final genome assembly is over 99.8% with properly alignments over 93.14%, an another individual of nearly 55 Gb of NGS reads were aligned back to the finally assembled genome using BWA (version 0.7.18) MEM^35^, achieving a 99.34% alignment rate, the properly paired alignments reached 86.53%.

## Data Availability

All sequencing data generated in this study are available in the China National Genomics Data Center under the accession number PRJCA051591. The genome assembly and annotation data are also available in the figshare website. All data will be made publicly available upon publication. Currently, the genome assembly result can be accessed upon request to the first author or corresponding authors.

## Code Availability

No custom scripts or codes were used during this study. All software used in this work is in the public domain, with parameters being clearly described in Methods. If no detail parameters were mentioned for a software, default parameters were used as suggested by developer.

### Acknowledgements

We are grateful to the Daxilin Forestry Farm for providing the samples

## Funding

This project was funded by the Special Project on Agricultural Financial Funding from the Ministry of Agriculture and Rural Affairs of China, entitled “National Special Project for the Precise Identification of Key Aquatic Animal Breeding Resources for Aquaculture”; the Heilongjiang Province Seed Industry Innovation and Development Fund Project, entitled “Identification, Evaluation and Breeding of Characteristic Aquatic Animal Germplasm Resources in Heilongjiang Province”; and the Government purchase for public service contract from Ministry of Agriculture and Rural Affairs of China (No.17230180), all to Guo Hu. This work was also supported by the Heilongjiang Postdoctoral Research Start-up Grant to Yuanyuan Zhang.

## Author contributions

Y.Y.Z., D.W., R.Z., S.W.L., X.H.Z. and G.H. conceived and designed the experiments; R.Z. conducted the sample collection experiments; Y.Y.Z. performed the genome assembly, repeat elements detection, gene homologues and genomic synteny analysis and the gene annotation; R.Z. and G.H. recruited animal resources; Y.Y.Z. and G.H. wrote the paper; and all authors read, edited and approved the final manuscript.

## Competing interests

The authors declare no competing interests.

---

**Supplementary Table 1.**
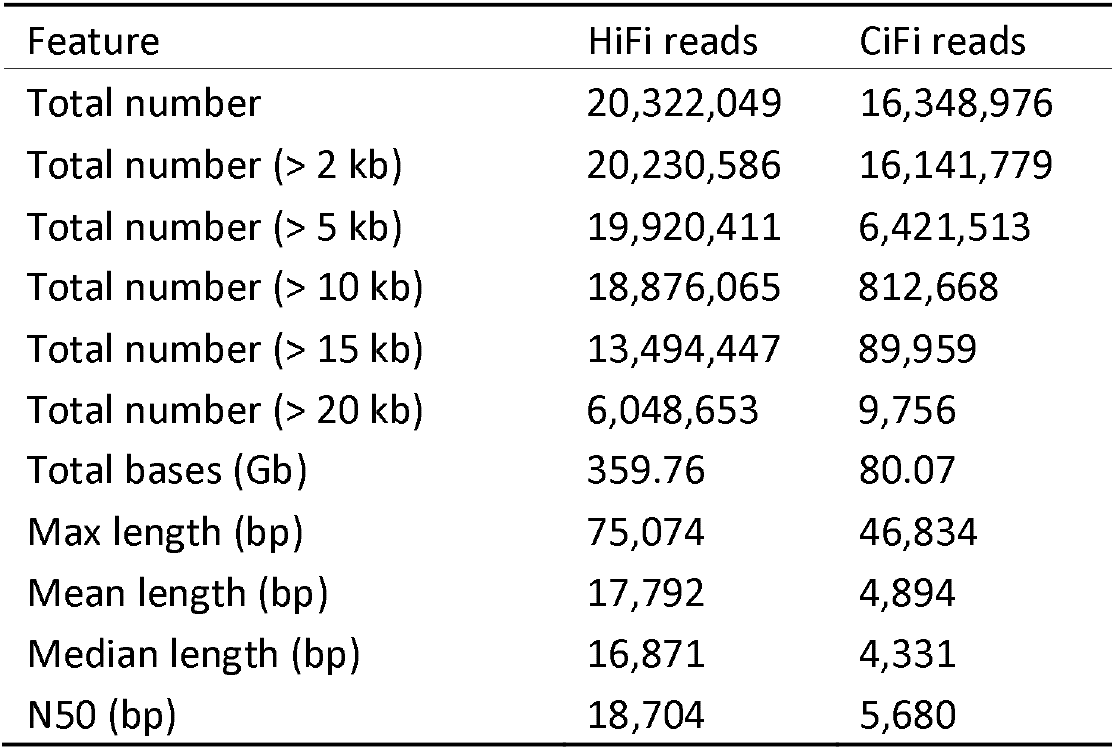
HiFi and CiFi reads of R. dybowskii obtained using the PacBio Revio platform.

**Supplementary Table 2.** The statistics of whole genome re-sequencing paired-end reads.

| SampleName | tissue | sex | ReadNum(M) | BaseNum(G) | GC(%) | Q30 (%) | Q20(%) |
| --- | --- | --- | --- | --- | --- | --- | --- |
| R1 | muscle | male | 1,947.45 | 292.12 | 44.17 | 95.71 | 98.56 |
| R2 | muscle | male | 183.05 | 54.91 | 43.21 | 95.47 | 97.10 |

**Supplementary Table 3.** The statistics of transcriptomic sequencing paired-end reads.

| SampleName | individual | Tissue | ReadNum(M) | BaseNum(G) | GC(%) | Q30_read1(%) | Q30_read2(%) |
| --- | --- | --- | --- | --- | --- | --- | --- |
| R1E | R1 | eye | 44.4 | 6.66 | 46.78 | 98.47 | 98.22 |
| R1G | R1 | testis | 70.28 | 10.54 | 41.03 | 97.47 | 95.53 |
| R1M | R1 | muscle | 47.47 | 7.12 | 48.56 | 98.4 | 97.52 |
| R1S | R1 | skin | 52.66 | 7.9 | 46.53 | 98.37 | 97.65 |

## Notes

### Competing Interest Statement

The authors have declared no competing interest.

